# Cancer Risk and the Somatic Cell Lineage Tree

**DOI:** 10.1101/2020.07.13.201004

**Authors:** Imre Derényi, Márton C. Demeter, Gergely J. Szöllősi

## Abstract

All the cells of a multicellular organism are the product of cell divisions that trace out a single binary tree, the so-called cell lineage tree. Because cell divisions are accompanied by replication errors, the shape of the cell lineage tree is one of the key determinants of how somatic evolution, which can potentially lead to cancer, proceeds. Cancer initiation usually requires the accumulation of a certain number of driver mutations. By mapping the accumulation of driver mutations into a graph theoretical problem, we show that in leading order of the mutation rate the probability of collecting a given number of driver mutations depends only on the distribution of the lineage lengths (irrespective of any other details of the cell lineage tree), and we derive a simple analytical formula for this probability. Our results are crucial in understanding how natural selection can shape the cell lineage trees of multicellular organisms in order to reduce their lifetime risk of cancer. In particular, our results highlight the significance of the longest cell lineages. Our analytical formula also provides a tool to quantify cancer susceptibility in theoretical models of tissue development and maintenance, as well as for empirical data on cell linage trees.

**Significance Statement:** A series of cell divisions starting from a single cell produce and maintain tissues of multicellular organisms. Somatic evolution, including the development of cancer, takes place along the *cell lineage tree* traced out by these cell divisions. A fundamental question in cancer research is how the lifetime risk of cancer depends on the properties of an arbitrary cell lineage tree. Here we show that for small mutation rates (which is the case in reality) the distribution of the lineage lengths alone determines cancer risk, and that this risk can be described by a simple analytical formula. Our results have far-reaching implications not only for cancer research, but also for evolutionary biology in general.

---

Cancer is a disease of multicellular organisms, which occurs when a somatic cell, after going through a number of genetic and epigenetic changes, starts to proliferate uncontrollably (1). It was Armitage and Doll (2, 3) who noticed that the incidence of various types of cancers in humans grows as a power function of age, and proposed a multistage model of cancer, where the exponent of the power function increased by unity (which typically falls between 5 and 7) corresponds to the number of driver mutations required for cancer initiation. Although we now know much more about carcinogenesis, the fundamental concept of the accumulation of a few critical mutations remains unchallenged (4–6).

All the cells of an organism are derived from a single cell along a single binary tree (as demonstrated in Fig. 1(a)). The extant cells (i.e., the leaf nodes) of a snapshot of the tree taken at a given moment of time are the ones that were either present in the organism at that time or had been lost by then. The internal nodes correspond to the cells that had already gone through a cell division. Similar lineage trees can be drawn for tissues or parts of tissues with the root node being the founder cell of the tissue or tissue part.

**Fig. 1.**
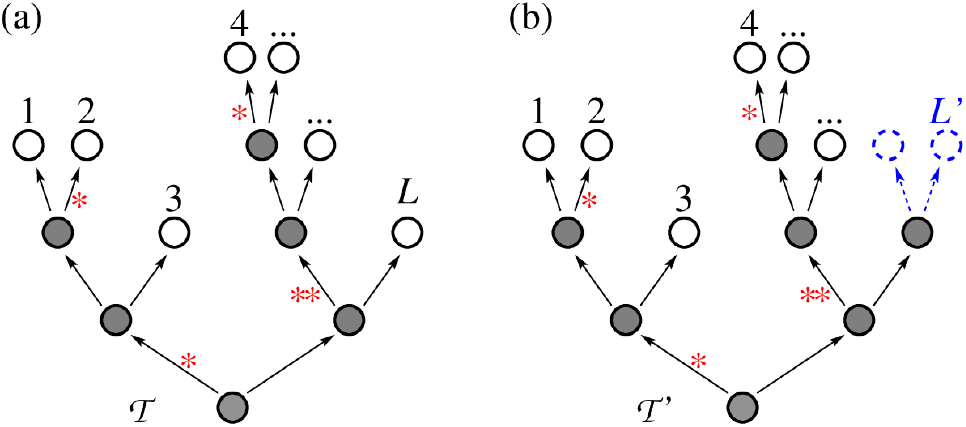
Illustration of a cell lineage tree with mutation accumulation. (a) A cell lineage tree 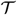 with *L* leaf nodes (white circles), and *L* − 1 internal nodes (gray circles). The lineage length (or divisional load) *D_i_* corresponding to leaf node *i* is the number of edges (cell divisions, denoted by arrows) leading from the root (bottom most) node to the leaf node. Mutations are indicated by red stars. (b) Cell lineage tree 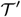 obtained by making leaf node *L* in tree 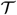 divide (indicated by blue dashed arrows and circles).

It is unclear (and might depend on the type of cancer) if individual driver mutations change the structure of the lineage tree or if they are neutral until a sufficient number is accumulated. The assumption of neutrality is consistent with the fact that the majority of cancers arise without a histologically discernible premalignant phase, and also with recent timing analyses, which suggest that driver mutations often precede diagnosis by many years, if not decades (7).

These observations indicate strong cooperation between driver mutations, suggesting that major histological changes that would significantly alter the structure of the lineage tree may not take place until the full repertoire of mutations is acquired (8).

Assuming that driver mutations occur at a uniform rate *μ* per cell division (indicated by red stars in Fig. 1), and that they are neutral until all the *m* required for cancer initiation are accumulated, we explore the fundamental question of how the structure of a cell lineage tree affects its risk of cancer.

## 1. Results

The above question can be translated into a mathematical (graph theoretical) problem: Given a binary tree 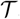, what is the probability 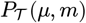 that a lineage (i.e., a path from the root node to a leaf node) with at least *m* driver mutations appears, if driver mutations are dropped to the edges uniformly with a frequency of *μ* per edge? In Fig. 1(a), e.g., the lineage belonging to leaf node 4 has 3 driver mutations (red stars).

Although no general analytical solution is known for this problem, an exact numerical answer can be given for any particular tree by following the procedure as outlined below. Every binary tree can be built up by subsequently merging its subtrees at their parent nodes, starting from the leaves. Each internal node (gray nodes in Fig. 1) is used for merging the two subtrees of its child nodes (daughter cells). The last merging at the root node of the entire lineage tree completes the merging sequence.

In a merging event let us denote the two sibling subtrees to be merged at their parent node by 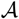 and 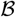. Let us denote the probability that the lineage with the largest number of driver mutations in subtree 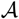 has exactly *j* driver mutations by 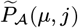, and in subtree 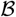 by 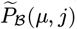. After extending both subtrees by adding the edge that leads to their common parent node to each of them, the probability that the lineage with the largest number of driver mutations has exactly *k* driver mutations in the such extended subtree 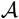 is

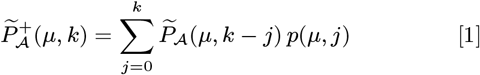

and that in the extended subtree 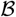 is

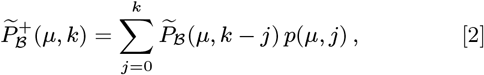

where

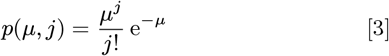

is the Poisson distribution with parameter *μ*.

After joining the two extended sibling subtrees at their parent node into subtree 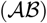, the probability that the lineage with the largest number of driver mutations in this newly merged subtree has exactly *m* driver mutations is

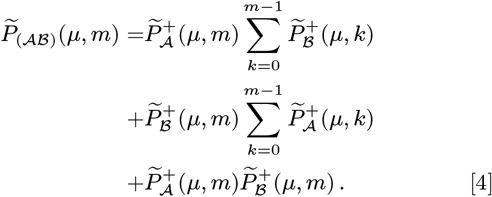

The initial condition for the merging sequence is that when the subtree (e.g., 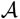) is a leaf node (i.e., a trivial graph consisting of a single node and no edges) 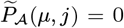 for *j* > 0 and 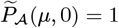.

After the merging process is completed the probability that the entire lineage tree 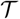 has a lineage with at least *m* driver mutations can be obtained as

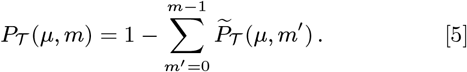

The reason that no closed formula exists for 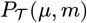 is the nonlinear nature of Eq. [4]. For small enough values of the driver mutation rate *μ* (such that 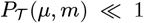), however, the dominant contributions of Eq. [4] arise when the summation factors (including the lineages with small numbers of mutations) are close to unity. Note that the very last term can be incorporated into either the first or the second summation. In this case Eqs. [1] to [4] become linear, and 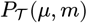 is expected to take the form

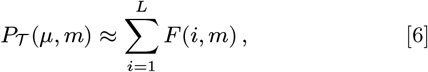

where index *i* runs over the leaf nodes from 1 to *L*. Furthermore, because the only quantity that is specific to a leaf node is its lineage length (or divisional load, denoted by *D_i_*), *F*(*i, m*) should depend on *i* only through *D_i_*. For small enough values of *μ*, *F*(*i,m*) is also expected to be dominated by its leading order term in *μ*, which is proportional to *μ^m^*, therefore, *F*(*i,m*) ≈ *μ^m^f*(*D_i_,m*).

Thus, for small enough driver mutation rates *μ* the risk of cancer (i.e., probability that a lineage with at least *m* driver mutations exists in the cell lineage tree) takes the form

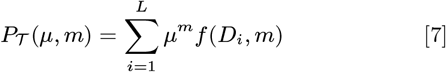

in leading order of *μ*.

The function *f*(*D_i_, m*) can be determined by introducing an additional division to the lineage tree (for which we chose leaf node *L*, as indicated by blue dashed lines in Fig. 1(b)). The probability for the new tree 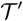 can be written up in two ways. First,

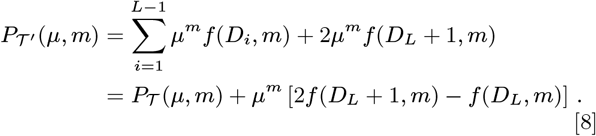

Second,

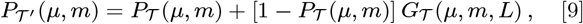

where the first term accounts for the possibility that a lineage with at least *m* mutations has already existed before adding the new division, and the second term corresponds to the scenario that no such lineage has existed, but with the elongations at least one of the two new lineages reaches the necessary number of driver mutations. The probability of this latter event is denoted by 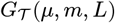. In leading order of *μ* the probability 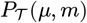 can be neglected from the second term and 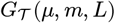 can be expressed as

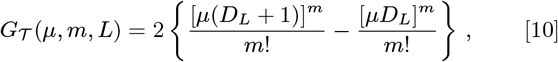

where the difference between the braces describes the excess probability conferred by a single elongation and the factor of 2 stands for the number of elongations. Combining the above, 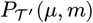 can be expressed as

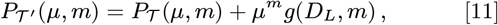

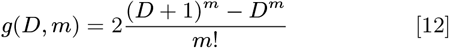

Comparing the two expressions [8] and [11] for 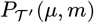 confirms the validity of our expectations on the form of 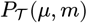 in Eq. [7] and leads to the recursive relationship

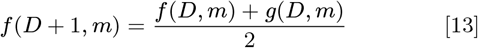

for the functions *f*(*D, m*), with the initial values of *f*(0, *m*) = 0 for any *m* > 0.

Expanding the recursion results in

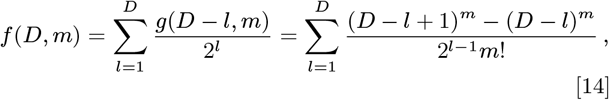

which cannot be simplified further. However, because in most real cell lineage trees the lineages (especially the longest ones that dominate the sum in Eq. [7]) are much longer than unity, *f*(*D,m*) can be well approximated by keeping only its highest order terms in *D*. For the two highest order terms

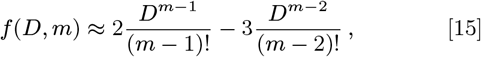

which is also equivalent to

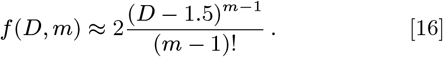

Plugging this approximation back into Eq. [7] leads to our main result that the risk of cancer, i.e., the probability of accumulating at least *m* driver mutations along a lineage can be well approximated by

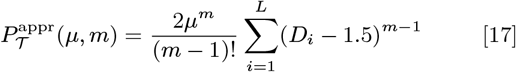

in leading order of *μ* and for the two highest orders of the lineage lengths. This approximation is very accurate for small enough mutation rates *μ* (such that 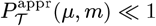, which is a valid assumption for the cancer risks of independent units of most human tissues), as demonstrated in Fig. 2 for a series of lineage trees that interpolate between the most skewed tree (linear chain with unit long side branches) and the most compact one (the perfect binary tree) with *L* = 2^16^ leaf nodes.

**Fig. 2.**
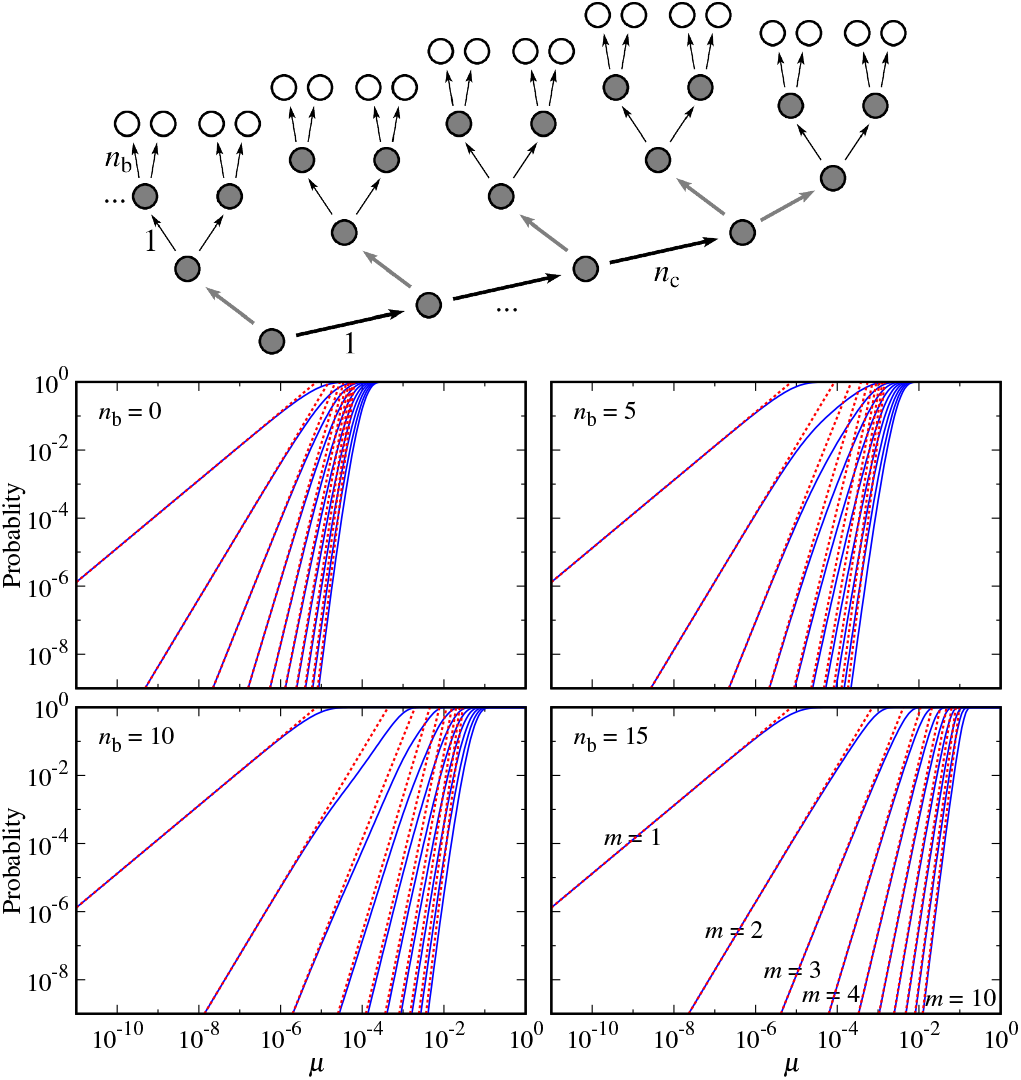
Examples for the probability of cancer in a series of cell lineage trees. The top panel illustrates how a series of lineage trees that interpolate between the most skewed tree (linear chain with unit long side branches, *n*_b_ = 0) and the most compact one (the perfect binary tree, *n*_c_ = 0) is generated: Identical perfect binary subtrees (indicated with thin arrows) of depths (lineage lengths) *n*_b_ are joined (with thick gray arrows) to an *n*_c_ long linear chain (indicated with thick black arrows). The number of leaf nodes can be expressed as *L =* (*n*_c_ + 2) 2^*n_b_*^. The four plots in the bottom panel show the exact probabilities 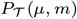 (blue lines, obtained form the merging process) that the lineage trees (for *L =* 2^16^ and for four different values of *n*_b_ = 0, 5, 10, and 15) have a lineage with at least *m* driver mutations (ranging between *m* =1 and 10 from left to right) as a function of the mutation rate *μ*, as well as their approximation 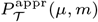 (red dashed lines) using Eq. 17. The approximation is very accurate as long as 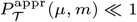.

## 2. Discussion

We derived our results for a cell lineage tree, where the root node is either the zygote or the founder cell of a tissue or a unit of tissue. Most tissue units (such as the colonic crypts), however, are sustained a set of stem cells, rather than a single one. Because our main formula [17] is a simple sum over all the leaves, it is also valid for a collection of trees, such as those generated by an initial set of stem cells.

We note that our results can also be applied to other evolutionary problems (e.g., maintaining cell cultures) where the divisions of the individuals are accompanied by mutations, and the accumulation of a given number of rare critical mutations has serious consequences.

The main message of our work is that assuming rare neutral driver mutations, the risk of cancer depends only on the distribution of the lineage lengths (irrespective of any other details of the lineage tree) and can be accurately approximated by a simple analytical formula. Because this formula is a sum of relatively high powers of the lineage lengths it is dominated by the longest lineages, which quantitatively explains why the minimization of the longest lineages (e.g., through hierarchical differentiation (9)) is crucial in minimizing cancer risk and somatic evolution in general.

Our results imply that the cumulative cancer incidence for any particular tissue should grow as a power function of age with exponent *m*, if the lengths of its longest lineages grow linearly in time. Indeed, for most large self-renewing tissues stem cells are known to divide at least several times a year (e.g., every 25 to 50 weeks (10) or every 2 to 20 months (11) for blood, and every 4 days (12, 13) for the colon) and, therefore, are expected to produce dominant linearly growing lineages.

We foresee applications of our method as more extensive cell linage tracing data becomes available, both at the organism (14, 15) and the tissue level (16).

## ACKNOWLEDGMENTS

The authors are thankful to Ágnes Backhausz for her valuable comments and advice. GJSz received funding from the European Research Council under the European Union’s Horizon 2020 research and innovation programme under grant agreement no. 714774 and the grant GINOP-2.3.2.-15-2016-00057.

